# Causal Evidence for the Neural Underpinnings of Subjective Happiness

**DOI:** 10.64898/2026.03.06.710111

**Authors:** Daniele Spica, Bertrand Beffara, Shira Cohen-Zimerman, Alexa Vushaj, Irene Cristofori, Jordan Grafman

**Author notes:** these authors equally contributed (Daniele Spica, Bertrand Beffara). Note: Questions concerning the Vietnam Head Injury Study can be directed to Dr. Jordan Grafman.

## Abstract

Happiness is a central but poorly localized dimension of human experience, and causal evidence linking discrete brain regions to subjective happiness is scarce. We tested whether focal brain damage modulates self-reported happiness by measuring Subjective Happiness Scale (SHS) scores in 131 male veterans with penetrating traumatic brain injury (pTBI) and 33 matched healthy controls (HC), and by applying voxel-based lesion–symptom mapping within anatomically pre-defined brain regions. Overall, individuals with pTBI reported higher SHS scores than HC. VLSM identified two lesion clusters associated with *increased* happiness after injury: a small cluster in the right anterior cingulate cortex and a larger cluster in the right orbitofrontal cortex. These results provide causal evidence that right frontal circuitry—notably the ACC and OFC—modulates subjective happiness, challenging current accounts of motivational and emotional processing and pointing to targeted neural substrates for understanding and potentially modulating human happiness.

## Introduction

Happiness can be viewed as the “ultimate goal” in life: when met, it signifies a life well-lived. In the scientific literature, happiness is often defined by subjective well-being (SWB), composed of both affective and cognitive well-being, which pertain to the balance between pleasure and displeasure in one’s life and the comparative judgement of one’s life against another, respectively (Andrews & McKennell, 1980; Schimmack et al., 2008). It is this subjective comparison, together with engagement in activities and goal achievement, that are believed to give rise to happiness (Lyubomirsky et al., 2005; Shin & Johnson, 1978). At first glance, happiness can be conceptualized as lying on one end of an affective continuum, where depressive states (McGreal & Joseph, 1993) lie on the other, with sadness situated in between these two ends (Tebeka et al., 2018, 2021). A corresponding line of reasoning would therefore result in targeting psychobiological correlates of sadness and/or depression (Chaudhury et al., 2015; Trapp et al., 2023) to alleviate them when medically required. However, despite advances in our understanding of depression, it still constitutes a large proportion of mental conditions (Goodwin et al., 2022; Ormel et al., 2022) and sadness is reported by an alarmingly substantial portion of the general population (Tebeka et al., 2018, 2021). As mentioned above, while treatment strategies have certainly focused on depression-related physiological systems (Chaudhury et al., 2015; Drobisz & Damborská, 2019; Frandsen et al., 2024; Kisely et al., 2018; Mayberg et al., 2005), the cognitive neuroscience literature on *happiness* is sparse. For example, the canonical relationship between happiness and sadness or depression has been questioned (Rafaeli & Revelle, 2006) and much less attention has been drawn to the psychobiological bases of happiness. The result is fewer and heterogeneous findings (Kringelbach & Berridge, 2009, 2011; Machado & Cantilino, 2016), thus yielding no satisfying agreed-upon cognitive neuroscience models of happiness (King, 2019).

In the current study, we adopted a hypothesis-driven approach to causally link brain lesions and happiness in human males. First, we aimed to identify relevant anatomical regions of interest (ROIs) to be used in our analysis. To this end, we reviewed existing theoretical works on the brain correlates of happiness *via* the PubMed database, using the following keywords associations: Happiness AND Brain; Happiness AND Neur*; Subjective well-being AND Brain; Subjective well-being AND Neur*. The search resulted in 3 review articles (Alexander et al., 2021; Kringelbach & Berridge, 2009; Suardi et al., 2016), with only one that reported clear brain activation patterns for each original article included in the review (Suardi et al., 2016). The works selected for this review (Suardi et al., 2016) involved “autobiographical recall” methods (i.e., they all elicited recall-based emotions) during brain imaging. Overall, recalls eliciting happiness (vs. neutral condition) activated the prefrontal cortex (PFC), the anterior cingulate cortex (ACC), the basal ganglia (BG), the insula, the midbrain, the temporal gyrus (TG) and the orbitofrontal cortex (OFC). Counterintuitively, some of these regions (i.e. PFC, ACC, insula, BG) were also found to be activated when contrasting recalls eliciting sadness vs. neutral conditions, as also suggested by a recent study that reported “antidepressant-like” effects under ACC inhibition in mice (Yuan et al., 2025). In reviewing the brain correlates of well-being, King (2019) concluded that the likeliest neuroanatomical correlates of well-being are located in the ACC. Together, these findings emphasize the lack of evidence and consistency regarding the brain correlates of happiness, and point to a general role of frontal structures in emotional processing.

One of the main caveats of human research on happiness is its non-causal approach, with studies primarily correlational in nature or concerning the effect of happiness on brain activity. A more straightforward way to causally investigate the brain correlates of happiness is the manipulation of brain functioning (Yuan et al., 2025). However, to our knowledge, no human study has employed this strategy. Another approach consists of assessing happiness in patients with disturbed brain functioning; still, there is almost no evidence of happiness following traumatic brain injury, except for one case report (Mendez & Parand, 2020), for which frontal lesions led to higher happiness-related traits. The current study aims to bridge this gap by measuring subjective happiness scores in a sample of patients with penetrating traumatic brain injuries and matched controls as part of the Vietnam Head Injury Study (Cristofori et al., 2024; Raymont et al., 2011). We adopted a confirmatory approach by defining anatomical regions of interest (ROIs), based on the existing literature reviews regarding happiness (King, 2019; Suardi et al., 2016) and applying Voxel-based Lesion Symptom Mapping (Bates et al., 2003). We predicted that lesions in the ACC, PFC, OFC, BG, TG, and insula would independently be associated with a modulation of happiness, as assessed using the Subjective Happiness Score (Lyubomirsky & Lepper, 1999).

## Methods

### Participants

Participants were adult male Vietnam veterans recruited from the Vietnam Head Injury Study (Raymont et al., 2011). We selected a total of 131 patients who suffered penetrating traumatic brain injuries (pTBI) together with 33 uninjured veteran healthy controls (HC) for a total of 164 participants. Lesion scans were acquired in Phase 3, approximately 35 years after the injury, at the National Naval Medical Center, Bethesda, MD, USA. Behavioral and neuropsychological data were collected during Phase 4, approximately 40-45 years post-injury. The CT scans administered during Phase 4 were considered stable, compared to earlier CT scans. The National Institute of Neurological Disorders and Stroke’s institutional review board approved the study protocols, and all participants gave written informed consent.

We compared pTBI and HC participants on a wide range of demographic and neuropsychological variables (see supplementary **Table S1**): age, years of education, handedness distribution, letter fluency total scaled score from the Delis–Kaplan Executive Function System (D-KEFS; Delis et al., 2001), total score from the Token Task (TT; Western Psychological Services, 1994), silhouette total raw score from the Visual Object and Space Perception Battery (VOSP; Thames Valley Test Company, 1991), and the total score from the Barratt Impulsiveness Scale (Patton et al., 1995). Additionally, the two groups were compared on the percentile score from the Armed Forces Qualification Test (United States Department of Defense, 1960), collected pre-injury and in Phase 4 for pre- and post-injury intelligence respectively.

### Subjective Happiness Scale

We used the Subjective Happiness Scale (Lyubomirsky & Lepper, 1999) as our measure of global happiness. The SHS is a questionnaire that measures a global and subjective judgement of the extent to which people feel happy or unhappy. Using 7-point Likert scales, participants had to answer 4 items, 2 of which require to describe themselves using absolute ratings and ratings relative to peers: (*“In general, I consider myself”: [“Not a very happy person”* to “*A very happy person*”] and “*Compared to most of my peers, I consider myself”: [“Less happy”* to “*More happy*”]) while in the remaining 2 items, participants had to judge how much the description of a happy and an unhappy person matches with them (*“Some people are generally very happy. They enjoy life regardless of what is going on, getting the most out of everything. To what extent does this characterization describe you?”: [“Not at all”* to “*A great deal*”]; and *“Some people are generally not very happy. Although they are not depressed, they never seem as happy as they might be. To what extent does this characterization describe you?”: [“Not at all”* to “*A great deal*”]). The average of the four items (with item 4 reverse-coded) yields a global subjective happiness score, with higher scores indicating greater happiness. For both controls and pTBI patients, as well as the whole sample, we assessed the internal consistency of the SHS using the Spearman-Brown formula. The Spearman-Brown method (Brown, 1910; De Vet et al., 2017; Spearman, 1910) first consists in splitting the items into two halves (i.e. odd items, 1 and 3 vs. even items, 2 and 4) and computing the score independently for both halves. Secondly, the scores from the two halves are entered into a correlation analysis. The correlation coefficient is then entered into the following formula: (2 x Correlation Coefficient)/(1 + Correlation Coefficient). For control participants, the pTBI patients and the whole sample, the Spearman-Brown index was 0.87, 0.80 and 0.82, respectively, indicating good internal consistency.

### Statistical Analyses

Statistical analyses for the comparison of SHS happiness scores between pTBI and healthy controls were performed under RStudio (PositTeam, 2025) using the *brms* package (Bürkner, 2017). A bayesian cumulative (“probit” link) ordinal regression mixed-effects model (Bürkner & Vuorre, 2019) was employed to assess the difference of SHS happiness scores between the two groups, assuming that the SHS reflects a continuous latent variable of happiness for each participant. The participant group was entered as the fixed effect, while the items and participants were entered as random effects. 1000 of the 3000 iterations were set as a warmup. Default (uninformative) priors were used. Control analyses were performed using Bayesian (categorical or canonical regressions where applicable) models under the *brms* package.

### CT scan acquisitions and normalization

Computerized tomography (CT) scans without contrast were obtained to identify brain lesions using a GE Medical Systems Light Speed Plus CT scanner at the Bethesda Naval Hospital in Bethesda, Maryland (USA). Magnetic resonance imaging (MRI) scans were not collected because of the likely presence of metallic fragments in our sample of participants with penetrating brain injuries. The reconstruction of the images was performed with the following parameters: in-plane voxel size of 0.4 × 0.4 mm, overlapping slice thickness of 2.5 mm, slice interval of 1 mm. Lesion locations and volume from CT images were acquired through the Analysis of Brain Lesion software (ABLe) run via the Medx medical imaging software version 3.44 (Medical Numerics, Germantown, MD) with enhancements to support the Automated Anatomical Labeling atlas (AAL, (Tzourio-Mazoyer et al., 2002). V.R. (a psychiatrist with clinical experience of reading CT scans) manually drew the lesions in native space on each 1 mm thick slice, and they were later reviewed by another researcher, J.G., who was blind to the results of the Phase 3 evaluation, enabling a consensus decision to be reached regarding the limits of each lesion. Scans were spatially normalized to the Montreal Neurological Institute MNI space (Collins et al., 1994) with the Automated Image Registration program (Woods et al., 1993) using a 12-parameter affine model on de-skulled CT scans.

### Regions of interest extraction

ROIs were selected according to the findings from Suardi et al. (2016), and masks were extracted from the AAL (Tzourio-Mazoyer et al., 2002) using SPM 12. For each mask, both areas in the left and right hemisphere were included, and a total of 6 masks were obtained: anterior cingulate cortex (corresponding to the left and right anterior cingulum, indexes 31-32), orbitofrontal cortex (corresponding to left and right olfactory areas, indexes 21-22; left and right orbital part of the medial frontal gyrus, indexes 25-26; left and right rectus, indexes 27-28; left and right orbital part of the superior frontal gyrus, indexes 5-6; left and right orbital part of the middle frontal gyrus, indexes 9-10; left and right orbital part of the inferior frontal gyrus, indexes 15-16), prefrontal cortex (corresponding to left and right superior frontal gyrus, indexes 3-4; left and right middle frontal gyrus, indexes 7-8; left and right pars opercularis of the inferior frontal gyrus, indexes 11-12; left and right pars triangularis of the inferior frontal gyrus, indexes 13-14; left and right medial part of the superior frontal gyrus, indexes 23-24), insula (corresponding to left and right insula, indexes 29-30), temporal gyrus (corresponding to left and right Heschl’s gyrus, indexes 79-80; left and right superior temporal gyrus, indexes 81-82; left and right superior temporal pole, indexes 83-84; left and right middle temporal gyrus, indexes 85-86; left and right middle temporal pole, indexes 87-88; left and right inferior temporal gyrus, indexes 89-90), and basal ganglia (corresponding to left and right caudate, indexes 71-72; left and right putamen, indexes 73-74; left and right pallidum, indexes 75-76).

### Voxel-based Lesion Symptom Mapping analyses

Voxel-based lesion symptom mapping analyses (VLSM) were performed with the VLSM2 toolbox version 2.60 (Bates et al., 2003) on Matlab R2023b. To compare the scores between patients with and without a lesion in a specific voxel, a general linear model was fitted at each voxel level, resulting in statistical maps showing voxels of the brain significantly related to a higher SHS score after a lesion, based on the behavioral results (see below). A total of 131 participants were considered for all the VLSM analyses. Lesion coverage and power maps for each voxel were obtained (Figure 2). A total of 6 analyses were performed using masks of the following ROIs in the left and right hemisphere: anterior cingulate cortex, orbitofrontal cortex, prefrontal cortex, insula, temporal gyrus, and basal ganglia. For each analysis, only voxels damaged in at least 4 patients and with sufficient statistical power (i.e. ≥ 0.8, see Kimberg et al., 2007) were included, with the cluster defining threshold set to 0.05. Lesion size was considered as a covariate (Pustina & Mirman, 2022). We used 5000 permutations to adjust for multiple comparisons, employing a minimum cluster size approach. Since the VLSM package we used does not provide effect sizes, we approximated them by computing custom estimates at cluster peaks. To this end, we first coded for each pTBI patient if they had a lesion at the cluster-peak coordinates based on individual CT scans. Secondly, we performed a canonical Cohen’s d computation based on the SHS scores for patients having vs. patients *not* having a lesion at the cluster peak coordinate. Of note, these canonical calculations do not account for covariate effects.

## Results

### Subjective Happiness Scale score

The Bayesian mixed-effects ordinal model of SHS scores as a function of the participant groups yielded a higher SHS score in pTBI patients than in matched healthy controls (β = 0.67, 95% CI [0.18, 1.16]; see **Figure 1**), showing an overall increase in happiness in patients with penetrating traumatic brain injuries (5.05 ± 1.03) compared to controls (4.42 ± 1.33). The next subsection addresses whether a lesion in specific brain regions drove this overall effect.

**Figure 1.**
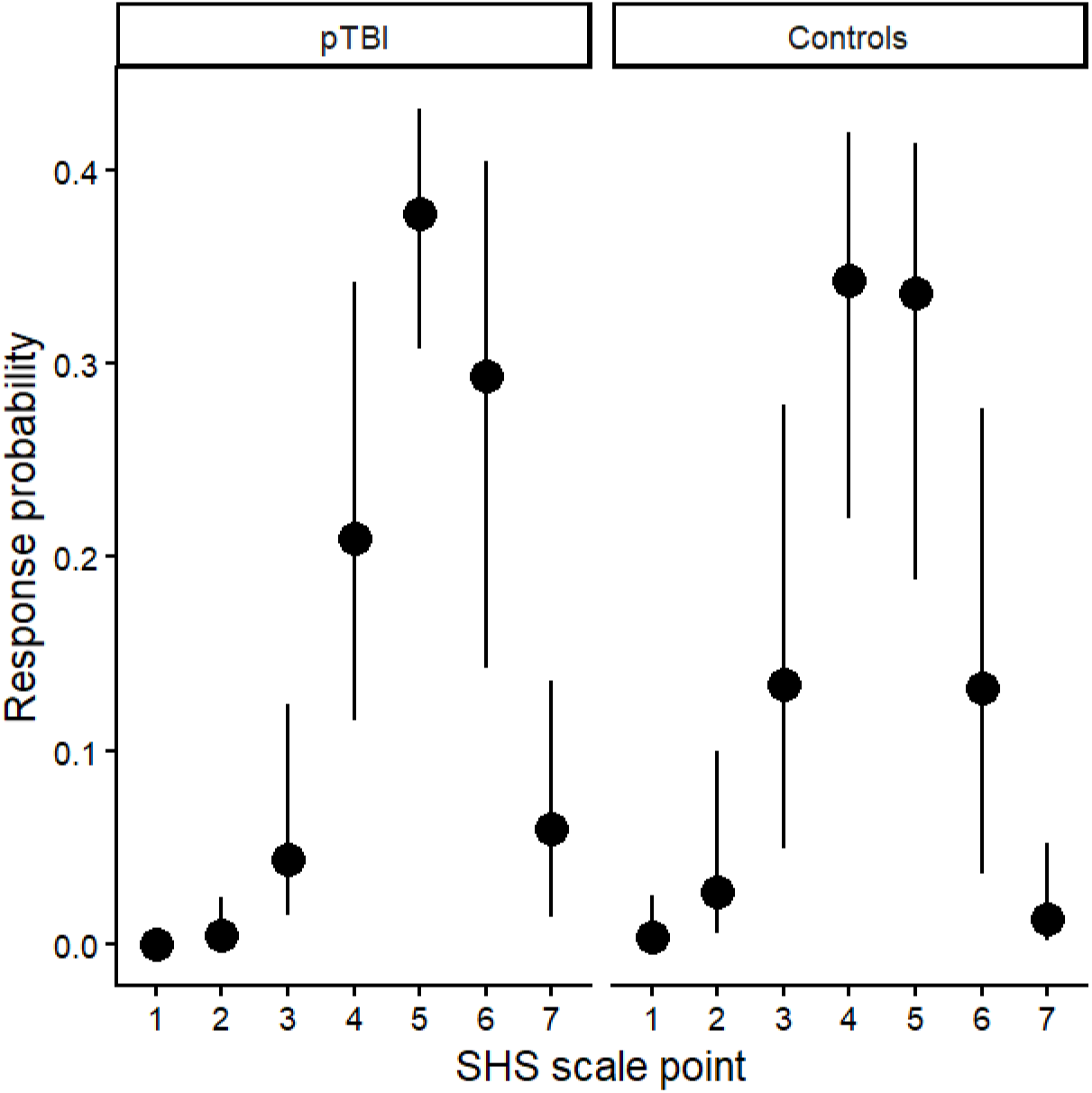
Ordinal regression estimates and the corresponding 95% credibility intervals for both pTBI and control groups for each SHS scale point. A score of 1 stands for lower happiness, while a score of 7 stands for higher happiness.

### Voxel-based Lesion Symptom Mapping

Lesions overlay and power maps are provided in **Figure 2a & 2b**, respectively. Of the 6 ROIs tested, only 2 remained significant after the permutation correction. We identified a significant cluster in the right ACC (**Figure 3**; volume = 14; MNI peak coordinates x = 16, y = 44, z = 12; MNI center of mass coordinates x = 16, y = 46, z = 11; Max T = 2.57; p = 0.0416; Cohen’s d computed from SHS scores in lesioned vs. other pTBI patients at MNI peak coordinates = 0.88) and a significant cluster in the right OFC (**Figure 4**; volume = 869; MNI peak coordinates x = 26, y = 50, z = -16; MNI center of mass coordinates x = 20, y = 54, z = -10; Max T = 3.42; p = 0.0498; Cohen’s d computed from SHS scores in lesioned vs. other pTBI patients at MNI peak coordinates = 1.15), which was located specifically in the orbital part of the middle, medial, and superior frontal gyrus.

**Figure 2:**
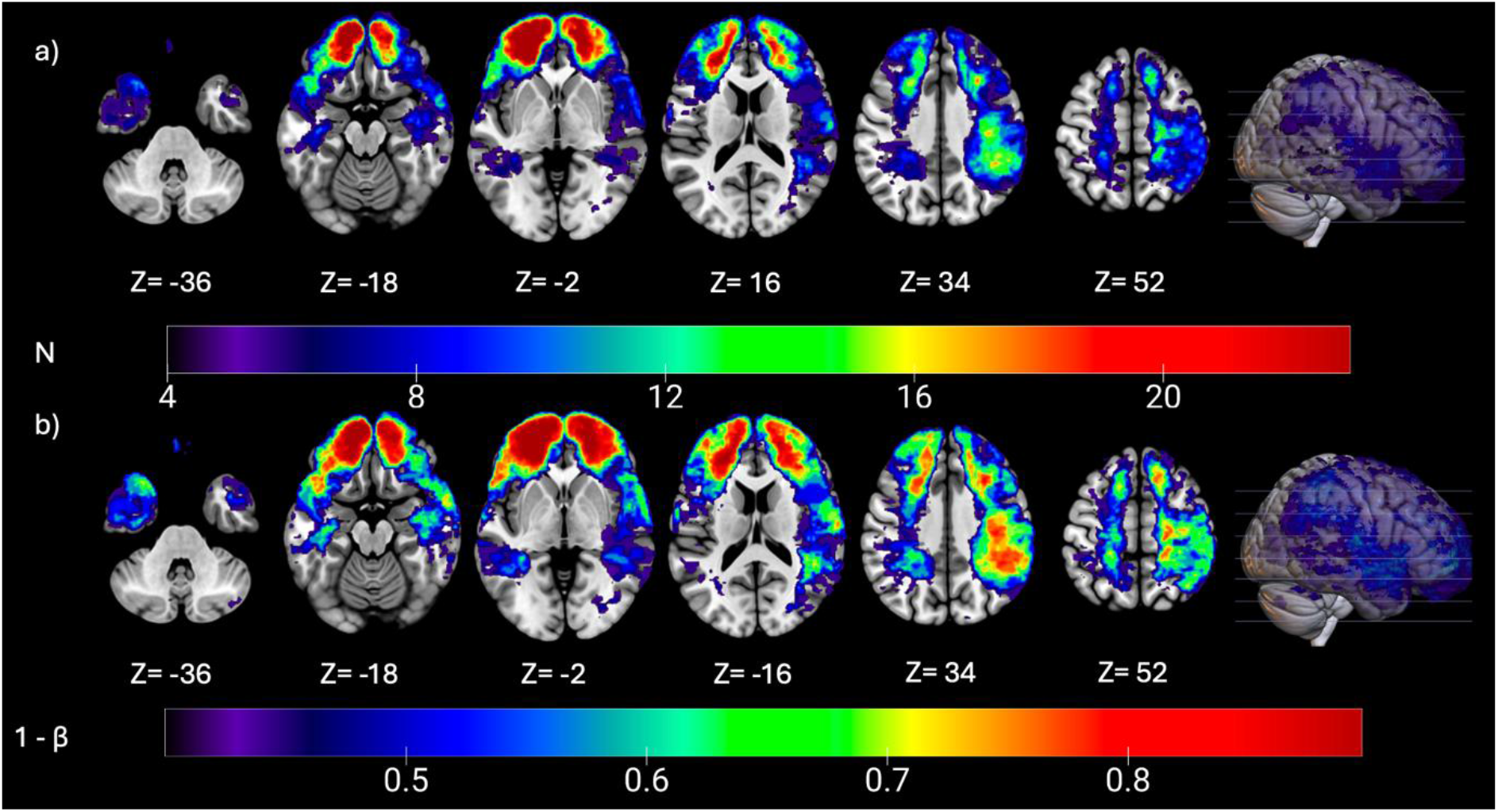
**a)** Lesions overlay for the 131 patients included in the VLSM analyses. The colorbar indicates the number of participants with a lesion at any given voxel. Only voxels damaged in at least 4 participants are shown. **b)** Power maps depicting the power distribution at any given voxel, for an alpha risk of 5% and an effect size of d = 0.8 (Baldo et al., 2010). Only voxels with at least 0.4 power are shown. We provide power descriptives in **Table S2**.

**Figure 3:**
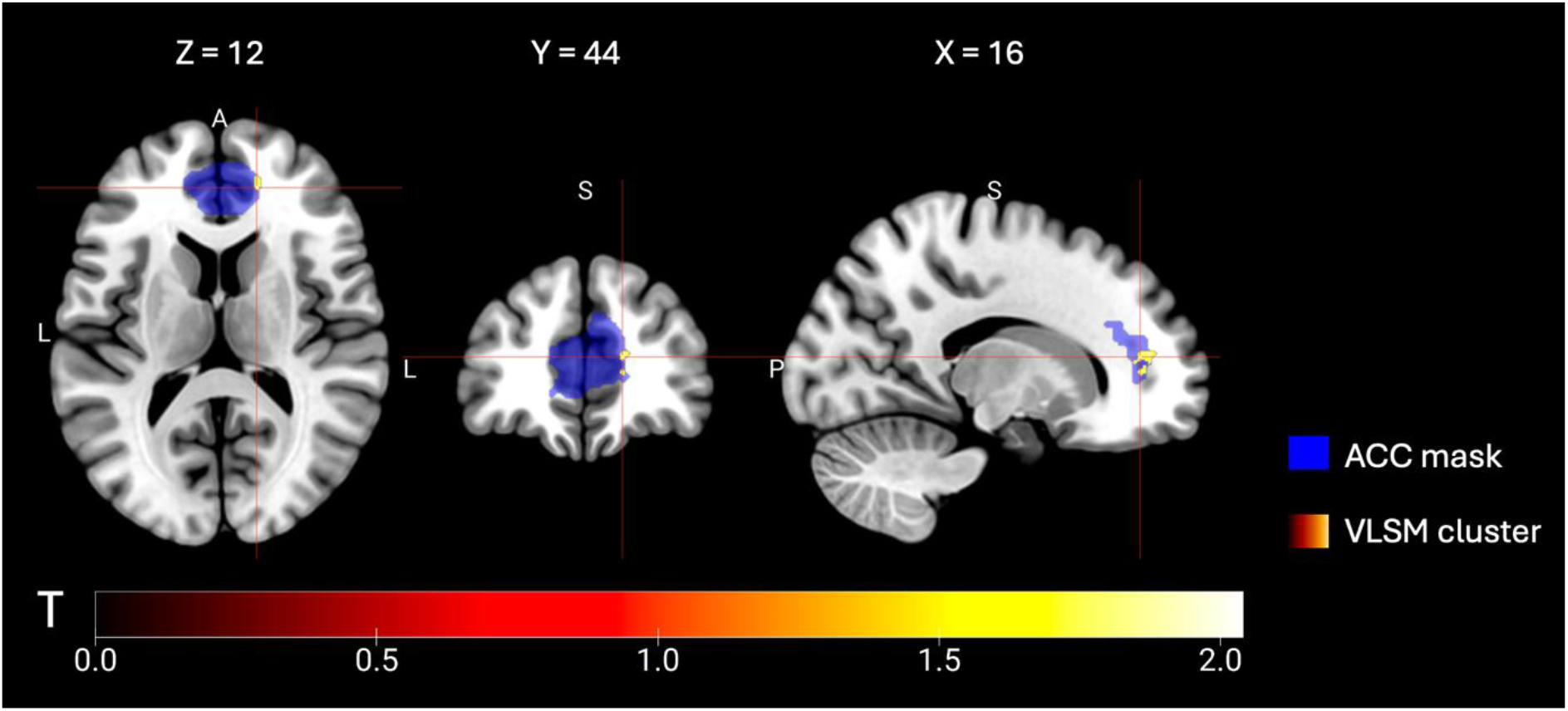
Anterior cingulate cortex (ACC) ROI from the AAL atlas (in blue) used for the VLSM analysis and significant ACC cluster (in pink/yellow). The colorbar indicates the T scores at any given voxel of the cluster. MNI coordinates: x = 16, y = 44, z = 12.

**Figure 4:**
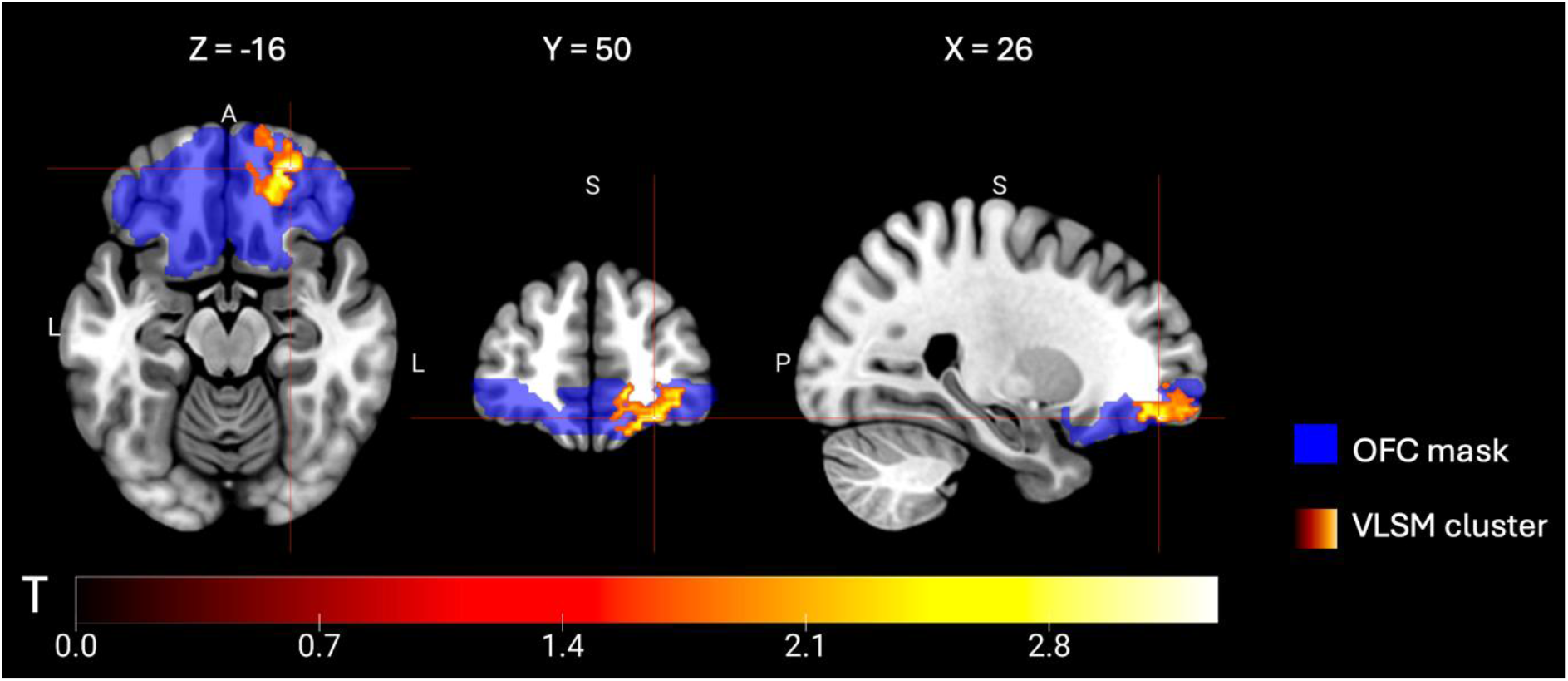
Orbitofrontal cortex (OFC) ROI from the AAL atlas (in blue) used for the VLSM analysis and significant OFC cluster (in pink/yellow). The colorbar indicates the T scores at any given voxel of the cluster. MNI coordinates: x = 26, y = 50, z = -16.

## Discussion

In this study, we used a Voxel-based Lesion Symptom Mapping approach to investigate the neural networks modulating happiness in a confirmatory way. On the behavioral level, pTBI patients reported greater happiness than matched healthy Vietnam combat controls. Based on pre-defined regions of interest, we found that lesions in the right anterior cingulate cortex and right orbitofrontal cortex were associated with a higher level of happiness.

The results of this study confirm that happiness, as an abstract idea, emerges from distributed brain computations. Suardi and colleagues (2016) offered an overview of the brain correlates of happiness and identified a restricted set of regions more consistently associated with well-being (i.e. frontal regions, plus limited evidence for reward and limbic systems and parietal and temporal regions *inter alia*; see also King, 2019, for similar patterns of results regarding well-being). While our study confirms frontal regions as brain correlates of happiness, additionally, we identified *causal* mechanisms for the emergence of human happiness: lesions in the anterior cingulate and the orbitofrontal cortices appear to modulate the subjective experience of happiness (see also Yuan et al., 2025, for complementary results concerning depression in mice; see Matsunaga et al., 2016). As previously mentioned, with the additional evidence we provided here, it is now clear that frontal regions contribute to the emergence of happiness. What is also theoretically quite sensible is the fact that these regions are certainly not *sufficient*. Hence, we will discuss the following below: 1) *how* the functioning of the orbitofrontal and the anterior cingulate cortices affects happiness and 2) what could be a typical neural circuit - with defined computational roles - of happiness.

A unique feature of cognitive science is that it can attempt to decompose higher-order functions into a series of local, basic, low-level mental processes. In an MRI study, Matsunaga et al. (2016) showed that the rostral ACC gray matter density was positively associated with subjective happiness and that experimentally inducing happiness activated the rostral ACC (see also Tanzer & Weyandt, 2020 for a meta-analysis of the brain correlates of happiness). They discussed these results first in light of autonomic regulation of emotional behavior, and secondly, in terms of outcome knowledge/reward estimation (see also Rolls, 2023, for a similar interpretation), calling for a role of the rostral ACC in the emergence of happiness.

How, then, could ACC lesions result in greater happiness? It may ultimately depend on the qualitative nature of ACC activity, in the sense that dysfunctional activity could hypothetically disturb efficient goal outcome knowledge and the related reward estimation and/or emotional regulation. Some studies report ACC activity associated with “social pain” constructs. For example, Eisenberger et al. (2003) showed that ACC haemodynamic activity *positively* correlates with social distress. Further, Cristofori et al. (2013) found that ACC theta activity selectively increased during social exclusion. In a fMRI study, Yoshino et al. (2010) reported that the increased ACC activity associated with painful stimulations was further enhanced by the co-presentation of sad faces visual stimuli. In addition, further directional connectivity analyses revealed close functional connections between the amygdala and the ACC (see below) under sad-painful stimulation contexts. Tolomeo and colleagues (2016) studied treatment-resistant depression patients with anterior cingulotomy (mid-cingulate part) performing an emotion recognition task and a Stroop task; larger lesion volume was associated with lower negative emotion recognition (disgust, anger, fear) - but, surprisingly, not sadness (but see Nakajima et al., 2022) - as well as higher Stroop interference.

Combined with additional evidence that ACC activity is linked to depression (Mayberg et al., 2005; Yuan et al., 2025), these studies converge towards a 2-component ACC role: one is social, affective, and emotional; the other is cognitive. Importantly, these 2 components may not relate to separate ACC subdivisions, but instead to the anterior, or pregenual part of the ACC. On these grounds, there are two predominant explanations of the increased happiness from ACC-lesioned patients. The first one is that disturbing ACC function would lead to both cognitive impairments *and* an affective bias towards positive emotions and reduced social distress, with the latter being more powerfully associated with increased happiness than the former. The second one is that our sample is exclusively composed of war veterans who may be subject to affective and emotion-related differences (e.g. depression and impulsiveness) compared to other individuals, although our data reveal these to be in the normal range (see table S1). As ACC over-activity is one of the underpinnings of treatment-resistant depression (Mayberg et al., 2005), it is possible that this feature may be over-represented in our sample; therefore, ACC lesions may relieve depression-like symptoms. In any case, our findings are compatible with the view of Rolls (2023), who places the ACC as a key node of reward-related positive emotional processing.

The orbitofrontal cortex also appears as a key region of the Rolls (2023) emotion-motivation model. Our OFC results fuel the same controversy as our ACC results: how could lesions in a key region involved in emotional and motivational processes (Rolls et al., 2020) lead to increased happiness? There is evidence that damage in the OFC is associated with higher recognition scores of happy faces and lower recognition scores of sad faces (Nakajima et al., 2022). Again, while we hypothesize that OFC lesions should disturb motivational, emotional regulation and goal-related cognitive processes (Rolls, 2023; Rolls et al., 2020), there seem to exist a second component inherent to the ACC/OFC function that relates more to emotion recognition, which corroborates our results. Further, the theoretical synthesis from Rolls (2023) acts as an overarching support to our result in that it elegantly links both the OFC and ACC roles to motivation, emotion, and goal outcome learning, which are constructs similar to psychological correlates of happiness (Nix et al., 1999; Wang & Milyavskaya, 2020). However, these two regions are not located along a single brain pathway for responding to positive and negative stimuli. Instead, they may primarily be involved in reflective processes linking behaviors to outcomes. It is possible that alternative routes, primarily involving the amygdala and the reward system (that were not tested as regions of interest in this study, see below) may underlie the setting of implicit happiness-related behaviors. Relatedly, Rolls (2023) proposes that the amygdala is unlikely to be involved in reported, experienced, and declarative emotions in humans. On the contrary, disturbing the OFC-ACC regions should result in poorly declared emotions, therefore providing an explanation for why OFC and ACC lesions resulted in biased happiness self-reports in our study. For example, Trapp et al. (2023) showed that patients with OFC lesions reported less depression (see also Koenigs et al., 2008, who reported reduced post-traumatic stress disorder after amygdala and frontal lesions).

The current study does not come without limitations. First, most lesions in our dataset were located frontally, reducing the likelihood of unexpected or novel findings in other brain regions. In addition, although we planned to use the basal ganglia as an ROI, this region lacked sufficient power for proper testing. Other limitations concern the generalization of our results, given that our sample only includes male participants. Finally, standard power consideration in VLSM studies use d = 0.8 as a smallest effect size of interest (Baldo et al., 2010), which is set as the default option in the VLSM package we used (Bates et al., 2003). This corresponds to a large effect size, which is not always observed in neurocognitive studies. We performed related power and effect size controls that are presented in the Results section (and see Supplementary Materials). In addition, we estimated effect sizes that were consistent with the *a priori* power approach used in the VLSM analyses.

To conclude, while most recent accounts point to a pivotal role of the right anterior cingulate cortex and right orbitofrontal cortex in motivation and emotional processing, here we found that lesions in these regions cause a *higher* self-reported happiness. We discussed these findings in light of existing models linking motivation, emotion regulation, and goal-related behavior, as well as evidence from emotion recognition. This discussion led us to consider two main tracts for future research. First, lesions in these two regions may bias individuals toward positive emotions. Second, they may impair reported, experiences, or declarative emotion. Future studies will be able to test these hypotheses using implicit naturalistic behavioral measures (e.g. daily-life goal completion, etc) as markers of happiness, using various and heterogeneous emotional tasks to assess for a general association between emotional processing and the orbitofrontal and anterior cingulate cortices functions. Clarifying how the orbitofrontal and anterior cingulate cortices shape both the experience and expression of happiness will be essential for refining current models of well-being and could open new avenues for the treatment of mood-related disorders.

## Acknowledgments

We are grateful to the Vietnam veterans who participated in this study. We thank V. Raymont, S. Bonifant, B. Cheon, C. Ngo, A. Greathouse, K. Reding, and G. Tasick for testing and evaluating participants. We also acknowledge the National Naval Medical Center and the National Institute of Neurological Disorders and Stroke for their support and facilities. The views expressed here are those of the authors and do not reflect the official policy or position of the Department of the Navy, the Department of Defense, or the U.S. Government.

