## Supplementary for "Causal Evidence for the Neural Underpinnings of Subjective Happiness"

**-**

**Supplementary Material**

Daniele Spica^1,2#^, Bertrand Beffara^1,2#^, Shira Cohen-Zimerman^3,4^, Alexa Vushaj^3^, Irene Cristofori^1,2^, Jordan Grafman^3,4,5^^

^1^ Institute of Cognitive Sciences Marc Jeannerod CNRS, UMR 5229, France

^2^ Department of Human Biology, University of Lyon 1 Claude Bernard, France

^3^ Cognitive Neuroscience Laboratory, Brain Injury Research, Shirley Ryan AbilityLab, Chicago, IL, USA

^4^Department of Physical Medicine and Rehabilitation, Feinberg School of Medicine, Northwestern University, Chicago, IL, USA

^5^Departments of Neurology, Psychiatry, and Cognitive Neurology & Alzheimer’s Disease, Feinberg School of Medicine, Department of Psychology, Northwestern University, Chicago, IL, USA

\# these authors equally contributed

^^^Note: Questions concerning the Vietnam Head Injury Study can be directed to Dr. Jordan Grafman.

For completeness, here we provide further details concerning our analyses’ power and their relationship with the tests’ statistics.

First, we found statistically significant VLSM clusters only in the right brain hemisphere (see Figures 3 and 4). Here, we controlled whether it was due to an overall power bias in the right vs. left hemisphere. To do so, we compared the mean power from the power map using specific left and right ACC/OFC ROI anatomical masks. A bayesian comparison showed a likely power difference between left and right ACC, with a higher power difference in the right vs. left ROI (β = 0.03, 95% CI [0.01, 0.04]; see **Figure 2a**). The opposite pattern of results was observed in the right vs. left OFC (β = -0.10, 95% CI [-0.11, -0.09]; see **Figure 2c**). Overall, these results did not specifically support a power bias between brain hemispheres. Next, we reasoned that if the observed significant brain clusters were due to a power bias, then we should observe a positive association between voxels’ T-values within the significant clusters and voxels’ power. For the ACC, we observed no likely association of this type (β = -1.60, 95% CI [-6.34, 3.04]; see **Figure 2b**). For the OFC, we observed a likely association, but in the reverse direction from the one that was expected (β = -2.50, 95% CI [-3.13, -1.89]; see **Figure 2d**). Together, these results suggest that the main pattern of observations in the current study does not (specifically) emerge from power issues (see also Discussion section).

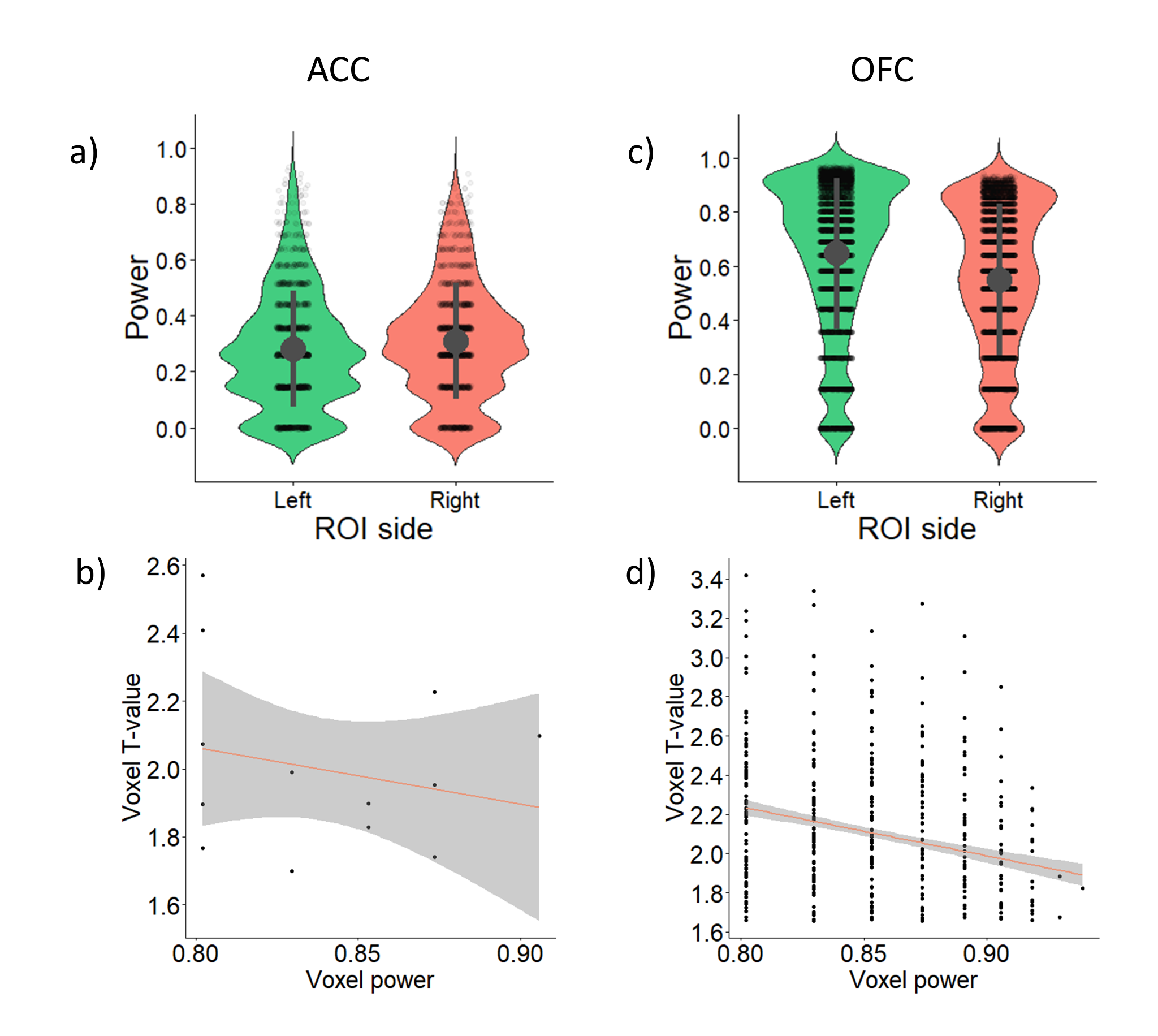

**Figure S1**: Power controls. a) Mean voxel power (± standard deviation) for each (left/right) side of the ACC ROI. b) Relationship between T-value for a given voxel and its power in the ACC ROI. c) Mean voxel power (± standard deviation) for each (left/right) side of the OFC ROI**.** d) Relationship between T-value for a given voxel and its power in the OFC ROI.

| **Measure** | **pTBI (n = 161)** | **HC (n = 33)** | **Slope**  **(pTBI vs. HC)** | **95% CI** | |
| --- | --- | --- | --- | --- | --- |
| Age | 63.30 (± 2.87) | 63.50 (± 3.76) | -0.19 | | [-1.39, 1.04] |
| Education | 14.55 (± 2.25) | 15.22 (± 2.20) | -0.63 | | [-1.49, 0.22] |
| Handedness^a^ | R: 107  A: 2  L: 22 | R: 25  A: 2  L: 5 | R: 0.03  A: -0.04  L: 0.02 | | R: [-0.11, 0.21]  A: [-0.16, 0.02]  L: [-0.14, 0.14] |
| Pre-injury intelligence^b^ | 64.58 (± 23.21) | 72.91 (± 17.06) | -8.34 | | [-18.93, 2.42] |
| Post-injury intelligence^b^ | 54.60 (± 26.08) | 70.06 (± 19.42) | -15.42 | | [-25.31, -5.67] |
| Verbal fluency | 8.77 (± 3.69) | 10.68 (± 3.74) | -1.91 | | [-3.37, -0.47] |
| Verbal comprehension | 96.85 (± 8.65) | 98.28 (± 2.02) | -1.43 | | [-4.56, 1.68] |
| Perceptual abilities | 19.93 (± 5.21) | 21.00 (± 3.64) | -1.07 | | [-2.95, 0.86] |
| Impulsiveness | 61.71 (± 10.23) | 62.87 (± 12.46) | -1.18 | | [-5.17, 2.96] |
| Depression | 7.83 (± 8.16) | 10.32 (± 8.89) | -2.49 | | [-5.82, 0.74] |

^a^R: right-handed; A: ambidextrous; L: left-handed.

^b^For healthy controls, “pre‑injury” and “post‑injury” refer to the two assessment time points matched to the TBI group’s timeline.

**Table S1:** Demographic and neuropsychological assessment of pTBI patients and controls.

|  | Mean ± standard deviation | Max | Min | % voxels ≥ 80% power |
| --- | --- | --- | --- | --- |
| OFC | 0.60 ± 0.29 | 0.97 | 0 | 32.99% |
| PFC | 0.52 ± 0.20 | 0.95 | 0 | 8.15% |
| ACC | 0.29 ± 0.21 | 0.93 | 0 | 1.88% |
| BG | 0.10 ± 0.11 | 0.52 | 0 | 0% |
| Insula | 0.40 ± 0.14 | 0.89 | 0 | 0.55% |
| TG | 0.43 ± 0.14 | 0.85 | 0 | 0.20% |

**Table S2:** Power descriptives for each ROI.
